# S373P Mutation Stabilizes the Receptor-binding Domain of Spike Protein in Omicron and Promotes Binding

**DOI:** 10.1101/2022.06.22.497114

**Authors:** Bin Zheng, Yuelong Xiao, Bei Tong, Yutong Mao, Rui Ge, Fang Tian, Xianchi Dong, Peng Zheng

**Affiliations:** State Key Laboratory of Coordination Chemistry, Chemistry and Biomedicine Innovation Center (ChemBIC), School of Chemistry and Chemical Engineering, Nanjing University, Nanjing, Jiangsu, 210023, China; Institute of Botany, Jiangsu Province and Chinese Academy of Sciences, Nanjing, 210014, China; State Key Laboratory of Pharmaceutical Biotechnology, School of Life Sciences, Nanjing University, Nanjing, 210023, China; Engineering Research Center of Protein and Peptide Medicine, Ministry of Education, China

## Abstract

A cluster of several newly occurring mutations on Omicron are found at the β-core region of spike protein’s receptor-binding domain (RBD), where mutation rarely happened before. Notably, the binding of SARS-CoV-2 to human receptor ACE2 via RBD happens in a dynamic airway environment, where mechanical force caused by coughing or sneezing occurs and applies to the proteins. Thus, we used atomic force microscopy-based single-molecule force spectroscopy (AFM-SMFS) to measure the stability of RBDs and found that the mechanical stability of Omicron RBD increased by ~20% compared with the wild-type. Molecular dynamics simulations revealed that Omicron RBD showed more hydrogen bonds in the β-core region due to the closing of the α-helical motif caused primarily by mutation S373P, which was further confirmed experimentally. Moreover, the binding ability of Omicron to ACE2 is promoted with a stabilized RBD. This work reveals the effect of the highly conserved mutation S373P which is present in most Omicron subvariants, including BA.1-5, BQ. 1, XBB, and CH.1.1.

## Introduction

The COVID-19 pandemic, caused by SARS-CoV-2, has been spreading worldwide for more than two years, primarily due to the continuous mutation of the virus^1,2^. As a single-stranded RNA virus that has infected a very large population, SARS-CoV-2 has undergone numerous mutations and rapidly adapts to the host. As the primary protein of the virus that binds to the angiotensin-converting enzyme 2 (ACE2) receptor of the host cell, the spike protein plays an essential role in the infectivity and transmissibility of the virus, showing an exceptionally high evolutionary rate (Fig. 1a)^3,4^. Mutations in the spike protein’s receptor-binding domain (RBD) are the most dangerous because this domain is the direct contact point with ACE2. In addition, the majority of neutralizing antibodies target the RBD^5–7^. Indeed, all SARS-CoV-2 variants of concern (VOCs) that have been announced by the WHO possess several mutations in the RBD^8,9^. These mutations lead to higher transmission, and these VOCs completely replace the original strain. Thus, knowledge of the effect of each mutation on the RBD is of great importance for understanding and fighting the virus.

**Fig. 1.**
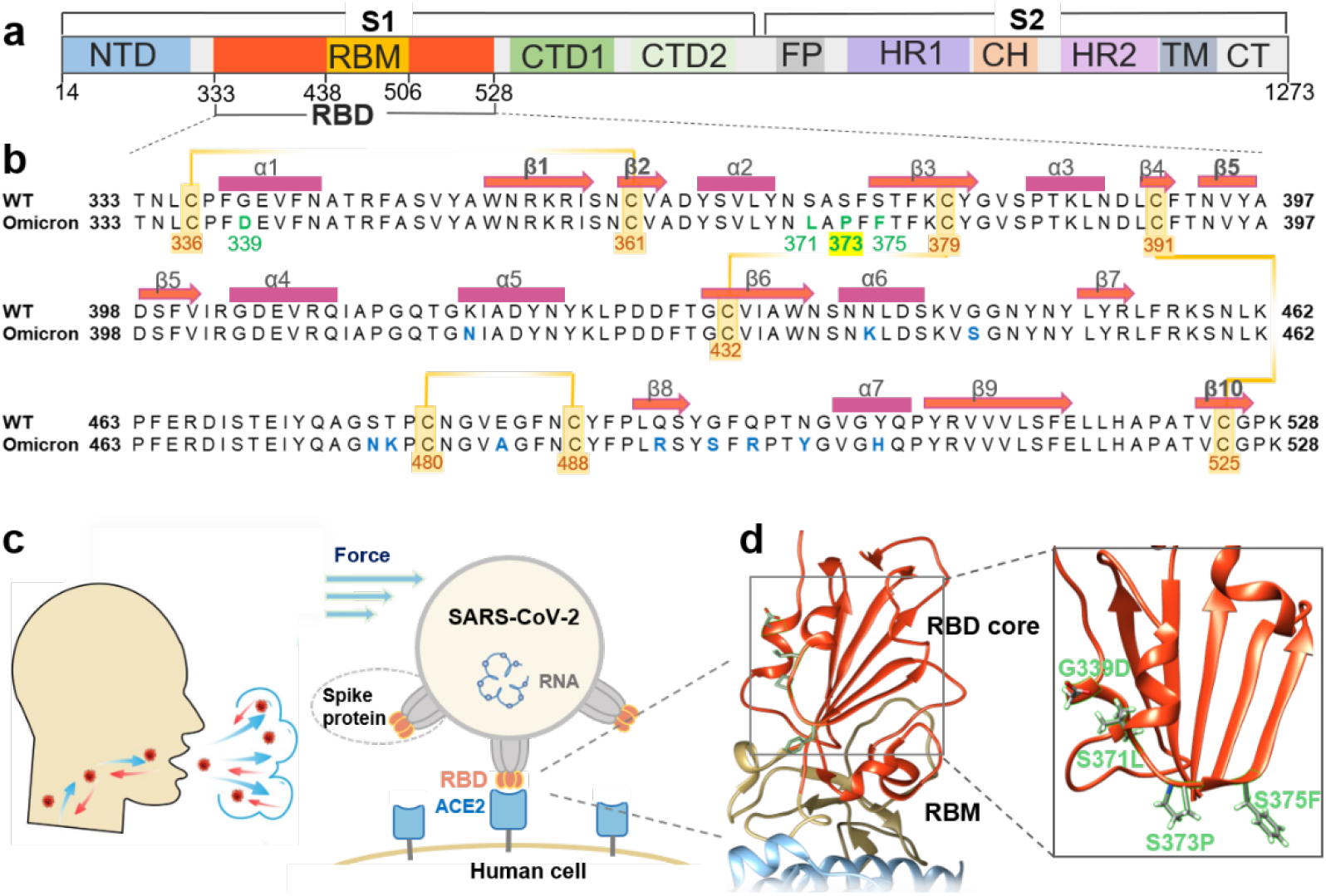
Newly occurring mutations at the β-core region of RBD in spike protein subjected to mechanical force. **a**, Domain architecture of the full-length SARS-CoV-2 spike (S) protein. **b**, Sequence alignment of the Omicron variant (BA.1) with the wild-type RBD (333-528). The mutations are highlighted in green or blue. The orange line indicates the four disulfide bonds with the residue number shown. The secondary structure elements of the RBD (PDB code: 6ZGE) are marked with rectangles (helices) and arrows (strands) lines above the sequence. **c**, The spike protein trimers (gray) on the SARS-CoV-2 virus particles could bind to ACE2 (light blue) on the human cell surface via the RBDs (orange). **d**, Crystal structure of the RBD:ACE2 complex (PDB code: 7wbl). The RBD β-core region and RBM are colored orange and yellow, respectively. Part of ACE2 is represented in light blue. Details of the Omicron variant structure are illustrated in the right panel. Four mutations that first appeared in Omicron are colored green.

The very recent VOC, Omicron (B.1.1.529), has more than 30 mutations in the spike protein, 15 of which are in the RBD, and shows the highest transmission thus far (Fig. 1a)^10–12^. Among these mutations, many with known effects are present on the flexible receptor-binding motif (RBM; residues 438-506), which makes direct interactions with the ACE2 receptor^13^. For example, mutations such as E484A and Q493R evade antibody neutralization, and Q498R and N501Y enhance spike/ACE2 binding. However, although containing so many mutations, the binding affinity of Omicron’s spike protein/RBD to ACE2 does not increase significantly, only similar to previous VOC Alpha and Beta. Thus, understanding the consequence of new mutations in Omicron RBD is still of great interest^14^.

A cluster of several new mutations, G339D, S371L, S373P, and S375F (Fig. 1b-c), is located close to the β-core region in the Omicron RBD (OmicronRBD)^10^. The effects of these mutations on the virus are still largely unknown. RBD has two subdomains with different stabilities and functions. The first is the flexible RBM with a concave surface for direct binding with ACE2, where mutations modify its binding affinity happen most. The second is a structured five-stranded antiparallel β-sheet core region that is covered with two connecting α-helices on both sides (Fig. 1c, colored in red). This region serves as a stably folded scaffold for the RBM, and these newly occurring mutations (green) in Omicron with undetermined functions are found in this mechanically stable region. In fact, as the critical component of the virus, the stability of the spike protein plays an important role in its entry efficiency and viral transmission^15^. Moreover, the binding of SARS-CoV-2 to the human host receptor ACE2 via the RBD happens in a dynamic airway environment, where mechanical force caused by coughing or sneezing occurs (Fig. 1c). Thus, viral attachment via RBD will be subject to the perturbations caused by mechanical force. Thus, we hypothesized that these new mutations modified the mechanical stability of Omicron, and a new strategy to study its stability is needed (Fig. 1d).

To this end, we used atomic force microscopy-based single-molecule force spectroscopy (AFM-SMFS) to investigate the mechanical stability of the RBDs from the wild-type (wtRBD) and Omicron variants. SMFS has been used to manipulate protein molecule(s) mechanically and studies the interaction between RBD/spike protein and its receptor in particular^16,17^. This technique mimics the external force from the dynamic airway on the protein during measurement, which the RBD may experience during binding to ACE2^18^. Previously, several groups have used SMFS to study the binding strength between the RBD and ACE2. They found that SARS-CoV-2 shows a higher binding force and binding energy to ACE2 than SARS-CoV-1^19,20^. We studied the effects of SARS-CoV-2 mutations and found that the N501Y mutation enhances RBD binding to ACE2, while K417N and E484K do not^21^. In addition to measuring protein-protein interaction, SMFS has been most widely used to measure the mechanical stability of the protein by (un)folding experiments^22–28^. Here, we used it to measure RBDs and found a significant force increase of Omicron compared to WT. Moreover, computational simulations were used to gain mechanistic insight into the mechanical unfolding process, and the results revealed the key mutations for the observed stability increment and the underlying mechanism.

## Results

We used a high-precision AFM-SMFS system to measure the stability of the RBD^29,30^. For an accurate comparison, the target protein RBD is fused with other marker proteins with known stabilities that are site-specifically immobilized in the system. Asparaginyl endopeptidase (AEP)-mediated protein ligation was used for covalent RBD immobilization on a peptide-coated glass coverslip, in which enzymatic ligation occurred between their specific N-terminal NH_2_-GL (Gly-Leu) dipeptide sequence and C-terminal NGL-COOH (Asp-Gly-Leu) tripeptide sequence (Fig. 2a, left panel)^31^. In addition, an ELP (Elastin-like polypeptide) linker with a defined length of ~50 nm was incorporated into the peptide, serving as a spacer to avoid the short-range nonspecific interaction. Finally, marker protein GB1 with a known unfolding force was used as an internal force caliper, and a dockerin-cohesin (Doc:Coh) non-covalent protein-protein interaction with a rupture force of ~500 pN was used as a single-molecule pulling handle^29,32^. Accordingly, the fused polyprotein Coh-(GB1)_2_-RBD-GB1-NGL was designed and immobilized for precise AFM measurements (Fig. 2a).

**Fig. 2.**
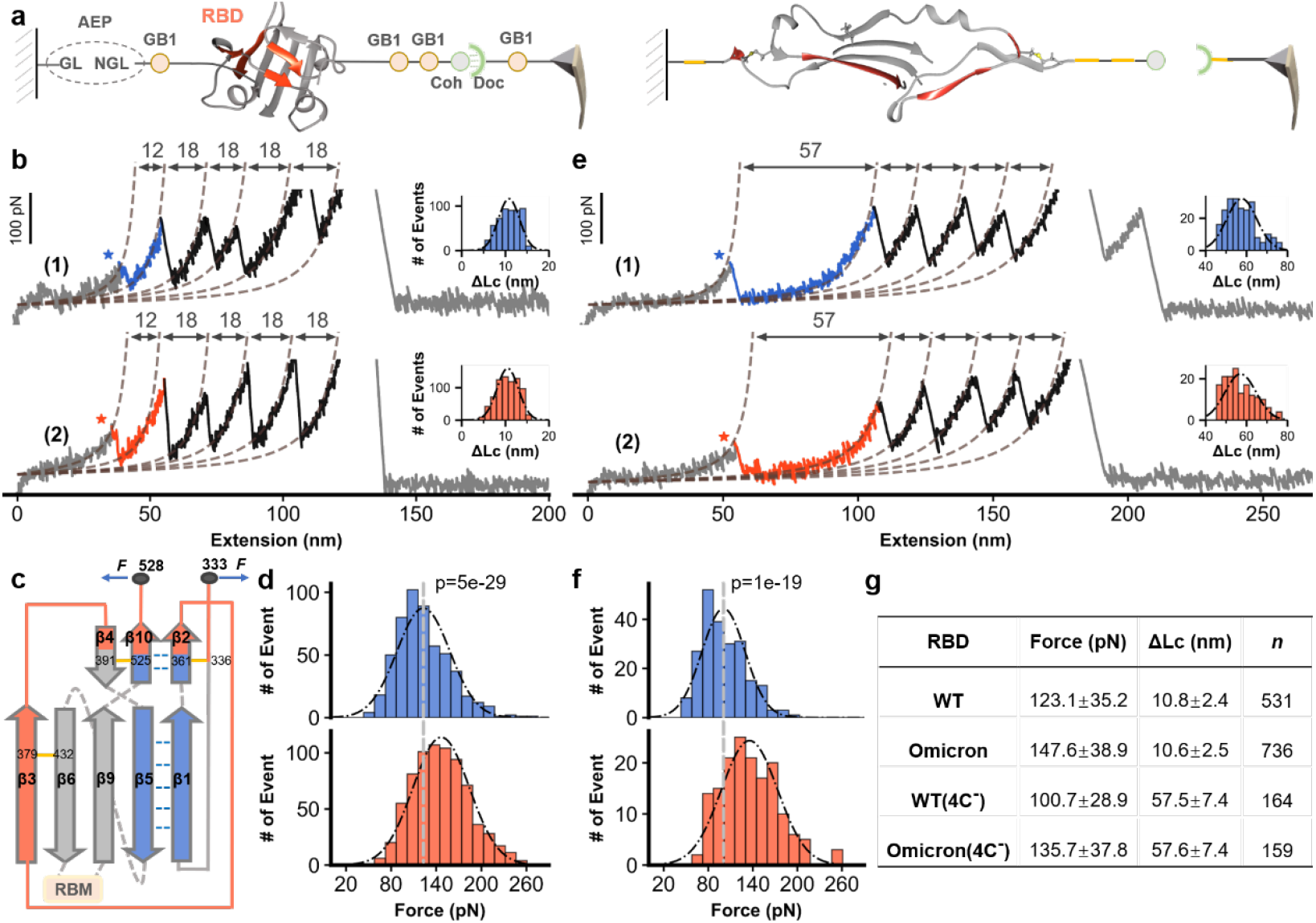
AFM unfolding experiment of wtRBD and OmicronRBD. **a**, High-precision AFM-SMFS system to measure the mechanical stability of the RBD using the fused protein Coh-(GB1)2-RBD-GB1-NGL. The protein was site-specifically immobilized to the glass surface using ligase AEP and could be picked up with a Doc-functionalized AFM tip via the Doc:Coh interaction (left panel). Then, the RBD and all other proteins were unfolded by stretching, and their mechanical stability was measured accordingly (right panel). **b**, Representative curves showing a series of sawtooth-like peaks from the unfolding of the RBD with a ΔLc of ~11 nm and marker protein GB1 with a ΔLc of ~18 nm. The Omicron and WT variants are colored orange and blue, respectively. The histogram of ΔLc is shown in the inset. **c**, The unfolded fragments are depicted by their secondary structures. **d**, The unfolding force of the OmicronRBD is ~20% higher than that of the wtRBD. P < 1e-5. P values were determined by two-sample t test. **e**, Unfolding of the RBD mutants with the deletion of two disulfide bonds (C391-C525, C379-C432) resulted in an elongated ΔLc from 12 nm to 57 nm, as expected. **f**, The force of the disulfide bonds-deleted Omicron RBD mutant was still higher than that of the WT. **g**, Detailed AFM data are shown.

Then, an AFM cantilever with a GB1-Doc-functionalized tip was used. By moving the tip toward the protein-immobilized surface, the RBD was picked up through the specific Doc:Coh interaction. Then, the tip retracted, and all of the proteins/domains sequentially stretched and unfolded, ending with the rupture of the strongest Doc:Coh interaction. The mechanical stability of the RBD has been measured accordingly (Fig. 2a, right). Finally, the tip moved to another spot on the surface and the cycle was repeated on different RBDs thousands of times. This system enables highly reliable and efficient AFM measurements and data analysis from a single molecule.

The fused protein containing wtRBD or OmicronRBD was built for AFM-SMFS measurements. Stretching led to a force-extension curve with an initial ~50 nm-length featureless curve from the extension of ELP linker, and then sawtooth-like peaks from the unfolding of GB1 (18 nm) and a peak from the rupture of the Doc:Coh interaction (Figs. 2b and S1). In addition to these auxiliary signals, a force peak with a ΔLc of ~11 nm was observed for both constructs, which suggests that the signal must result from RBD unfolding. Statistically, the value was 10.8±2.4 nm for the wtRBD and 10.6±2.5 nm for the OmicronRBD. To confirm these results, we calculated the theoretical number of extendable residues upon RBD unfolding. First, four disulfide bonds are present in the RBD, which cannot be broken, and the residues between them are locked^33^. Thus, RBD unfolding will lead to the extension of only 31 residues, from the first structured residue C361 to the last residue C391 (Fig. 2c, red). As a result, the theoretical ΔLc value is 10.3 nm (31*0.365 nm/aa-1.0 nm), agreeing well with the experimental value of ~11 nm and confirming that the peak was attributed to the RBD. Thus, the average unfolding force was 123.1 pN for wtRBD and 147.6 pN for OmicronRBD (Fig. 2d). These results support our hypothesis that the OmicronRBD has increased mechanical stability upon mutation.

To further verify that the observed data were from RBD unfolding, we deleted two disulfide bonds (C379-C432 and C391-C525) by mutating the four cysteines to alanines. As a result, 190 residues that the two disulfide bonds had locked in the RBD and previously inextensible will be unfolded in this construct, which led to a larger ΔLc of 58.1 nm (162*0.365 nm/aa-1.0 nm) (Supplementary Fig. 2). Then, we performed the same AFM experiments on these two constructs. As expected, the 11 nm peak mostly disappeared, while a new peak with a ΔLc of ~58 nm was observed (Fig. 2e), confirming that the previous 11 nm peak was from the RBD. These values were 57.5 nm for the WT and 57.6 nm for the Omicron. Moreover, the unfolding force of OmicronRBD (135.7 pN) was higher than that of wtRBD (100.7 pN) for this mutant (Fig. 2f, 2g), again demonstrating the stronger stability of OmicronRBD than wtRBD. To gain mechanistic insight into the considerable stability enhancement and pin down the exact mutation(s) responsible for this effect, we performed two different all-atom molecule dynamics simulations, focusing on the hydrogen bond in the β-core region of the RBD^34^. First, we performed steered molecular dynamics (SMD) simulations to visualize the mechanical unfolding process of the RBD and reveal the unfolding barrier/key force-bearing motif. Similar to the above experiment, the RBD was stretched with a constant pulling speed under a simulated force field and showed the force-extension trajectory (Fig. 3a, Video S1 for WT, Video S2 for Omicron). Indeed, the unfolding force of the OmicronRBD was generally higher than that of the wtRBD over the trajectory. This difference was quantitatively reflected by the potential of mean force analysis from irreversible pulling through Jarzynski’s equality (Fig. S3), which gave a value of 205 kJ/mol vs. 184 kJ/mol (Fig. 3a, inset). Moreover, the SMD trajectory demonstrated that the major contributing factors to the unfolding barrier are from the peeling between β-strands 2 and 10 and β-strands 1 and 5, as shown by snapshots of the three key unfolding events (Fig. 3a, red). Therefore, the interaction between these β-strands determined the mechanical stability of the RBD.

**Figure 3.**
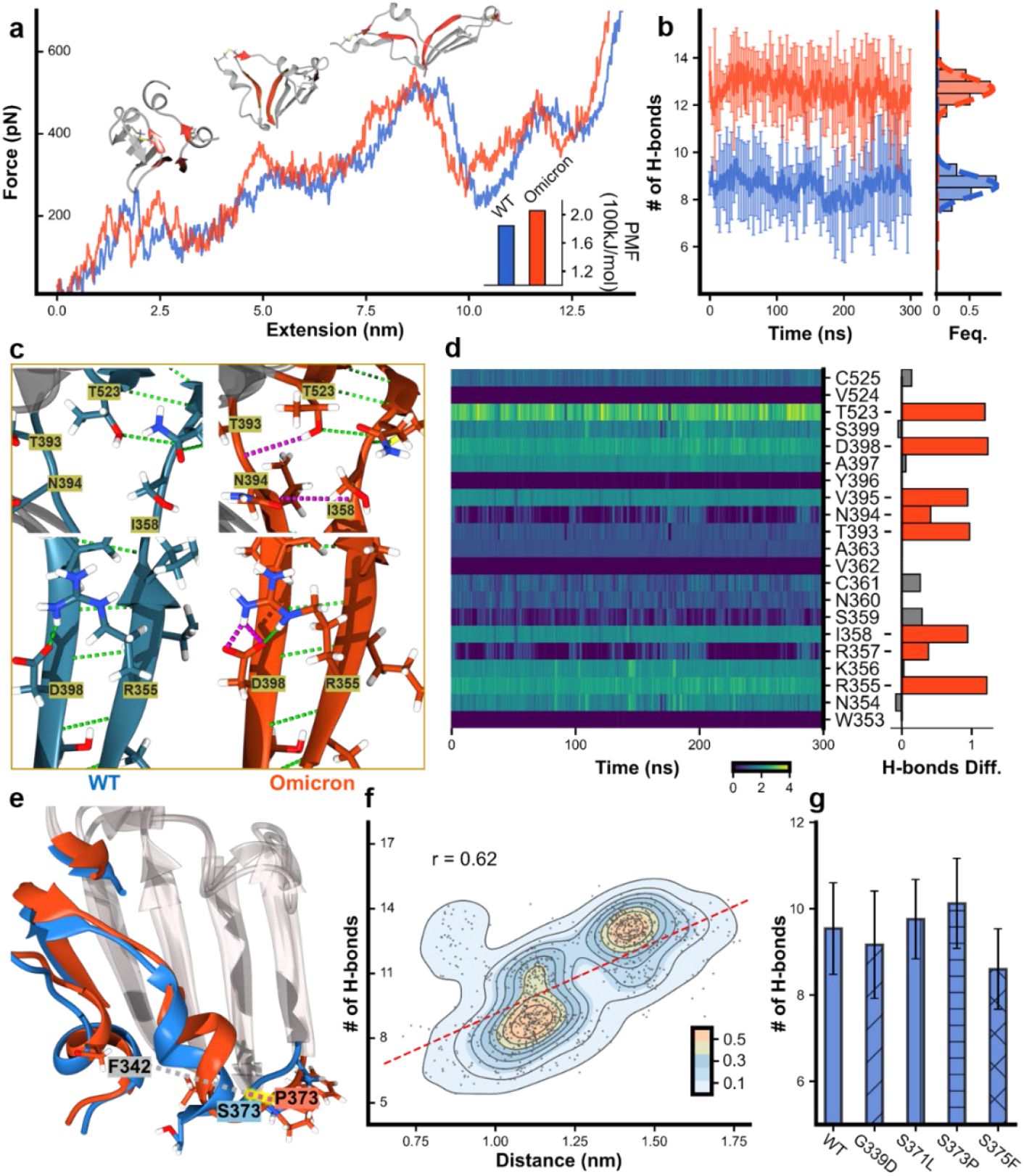
MD simulation results for wtRBD and OmicronRBD focusing on the four force-bearing β strands. **a**, SMD trajectories of stretching wtRBD (blue) and OmicronRBD (orange). The three key snapshots show the renderings of the RBDs at different times. The four force-bearing β-strands, β2/β10 and β1/β5, in the β-sheet are colored red. Inset: the potential of mean force (PMF) calculated. **b**, MD simulations show the number of hydrogen bonds formed in the four β-strands. **c**, The enlarged structure shows the hydrogen bonds (dashed line) in the four β-strands. Additional H-bonds in Omicron are colored pink. **d**, The evolution of the number of hydrogen bonds for each residue involved in β2/β10 and β1/β5 during the MD simulations. The difference in hydrogen bonds between Omicron and WT was shown in the right panel, and the red bars indicate a noticeable increment in Omicron compared to WT. **e**, Overlapped structure of Omicron and WT focusing on the β-core region of the RBD shows that the conformation of α2 in Omicron has changed. The distance between P373 and F342 in Omicron is longer (red dash line) than that between S373 and F342 in the WT. **f**, The relationship between the distances of residues 373 near α2 and F342 near α1 (as a reference point) and the number of H-bonds was extracted from multiple MD simulations. **g**, By introducing a single mutation into the wtRBD, MD simulations show that S371L and S373P increase the number of hydrogen bonds in the region.

Thus, we performed MD simulations to determine the number of hydrogen bonds in these four β-strands of the wtRBD (Video S3) and OmicronRBD (Video S4). After 300 *ns* of 10 times simulation, ~9 hydrogen bonds were present in the wtRBD, while ~13 bonds were found in the OmicronRBD, on average (Fig. 3b). Moreover, a detailed analysis of the hydrogen bond number for each residue along the simulation for both wtRBD and OmicronRBD was also performed, showing that several residues participated in more hydrogen bonds in the Omicron (Figs. 3c, 3d, S4 and S5). For the region of the force-bearing β-strands, we also analyzed the energy terms (Fig. S6). The short-range LJ (Lennard-Jones) potential has no obvious change between wtRBD and OmicronRBD. However, the Coulomb interaction of OmicronRBD is stronger than wtRBD. Thus, the Coulomb interaction might play an important role in the stability of the β-strands and can be affected by the sodium chloride in the solvent.

To rationalize the origin of this difference in the number of hydrogen bonds, we overlapped the structures of the WT and Omicron RBDs, focusing on the β-core region. Notably, α-helix 2 in Omicron moved toward the β-sheet core, as reflected in the cryo-electron microscopy (cryo-EM) structure (Supplementary Fig.7). Although this movement is not apparent in cryo-EM structure with a fixed conformation, it can be readily inspected through the structure from MD simulations with multiple dynamic conformations (Fig. 3e, highlighted in red). To validate the relationship between this structural change and the number of hydrogen bonds, we performed ten times of MD simulations (300 *ns*) and averaged the results. Then, we chose every six ns to calculate the number of hydrogen bonds in each simulation. Since the position of α-helix 1 is relatively consistent, we chose it as a reference point, and measured the distance between residue 373 in α-helix 2 and residue F342 in α-helix 1 at each section. These distances were plotted along with the number of hydrogen bonds. As shown in Fig. 3f, the longer the distance between them, the more hydrogen bonds are generally observed. The relationship between distance of residue 371/375 to residue F432 and the number of hydrogen bonds is not very significant (Supplementary Fig. 8).

The new mutations G339D, S371L, S373P, and S375P in the core region of the RBD are close to α-helix 2, suggesting their roles in this structural change. Thus, we built a single mutation into the wtRBD for MD simulations (*n*=10) and analyzed the corresponding number of hydrogen bonds. On average, wtRBD(S373P) showed the highest increase of two hydrogen bonds, and wtRBD(S371L) showed the modest addition of one hydrogen bond, and the effects for the other two mutations were trivial (Fig. 3g). Thus, it appears that the S373P mutation rigidifies the local polypeptide chain, leading to the inward movement of helix 2 from the structure. As shown in supplementary Figs. S9 and S10, the rigid proline 373 in OmicronRBD might open the loop between the nearby helix and β-strand, so that it can maintain a relatively large distance from the α-helix. Since the helix is directly connected to β1/β2, this conformational change may allow these β-strands to form more hydrogen bonds in a better orientation. Consequently, MD simulations revealed that S371L and S373P most likely account for the increased stability of the OmicronRBD.

Inspired by the simulation results, we built four wtRBD mutants in the polyprotein constructs, focusing on residues 371 and 373 for experimental confirmation. First, wtRBD(S371L) and wtRBD(S373P) were constructed, in which these two mutations found in Omicron were added to the wtRBD. As expected, the unfolding signal of the RBD was the same except for the force (Figs. 4a and S11). The AFM results for wtRBD(S371L) showed no force increment (123.4 pN), while wtRBD(S373P) showed a considerable increment of 20 pN (143.8 pN).

**Figure 4.**
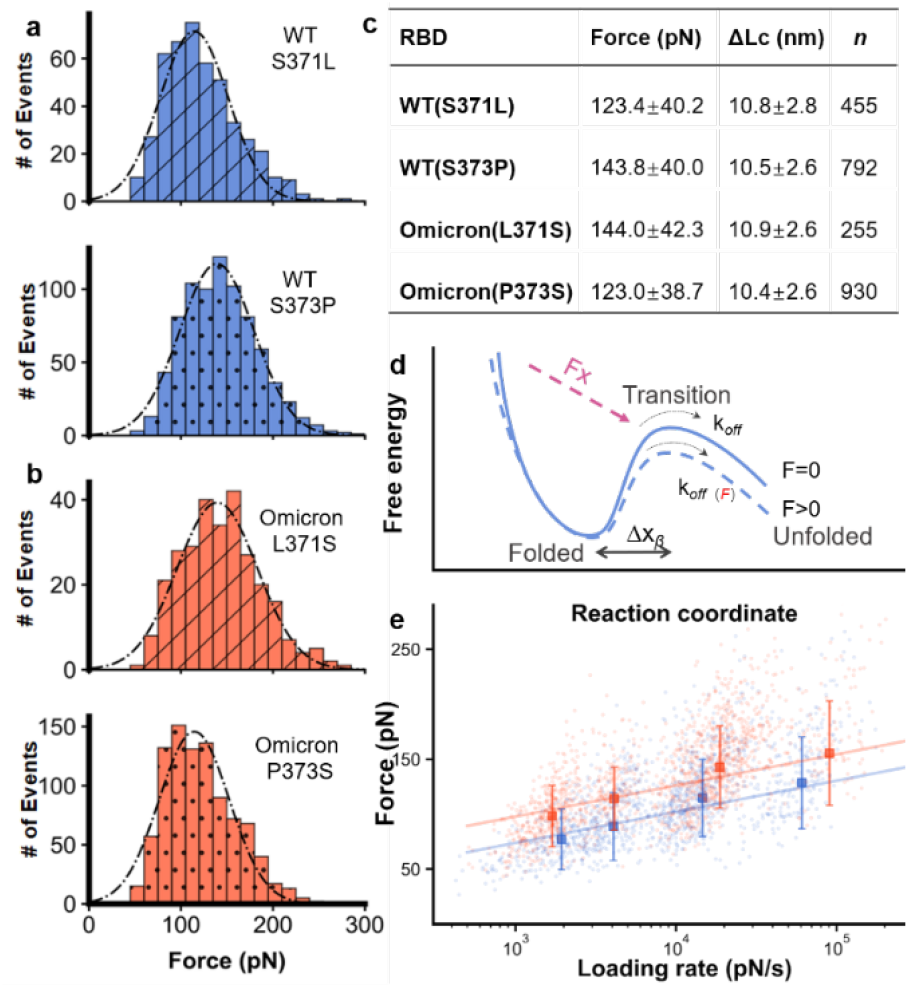
AFM-SMFS results of RBDs with a point mutation. **a-b**, Unfolding force histograms of wtRBD(S371L), wtRBD(S373P), OmicronRBD(L371S), and OmicronRBD(P373S) showing that residue 373 is the primary site for the fine-tuning of RBD stability with the most significant force change upon mutation. **c**, The unfolding force of RBDs is shown. **d**, The free-energy potential between folded state and transition state of the protein in the absence (solid, blue) and the presence of an externally applied pulling force (dashed, blue). **e**, The unfolding forces of the WT and Omicron RBDs show a linear relationship with the logarithm of the loading rate.

Moreover, we built OmicronRBD(L371S) and OmicronRBD(P373S), in which the mutation found in Omicron is reverted back to the residues present in the wtRBD while keeping all other mutations. The unfolding force decreased in these RBDs. The unfolding force of OmicronRBD(L371S) decreased slightly to 144.0 pN, while that of OmicronRBD(P373S) decreased to 123.0 pN (Fig. 4b-c). Also, we built a similar point mutation on the remaining two residues, 339 and 375, in the cluster and performed the same AFM experiments. Agreeing with the computational simulation, the mutation on these two sites did not change the mechanical stability of the RBD (Supplementary Fig. 12). Consequently, this experimental measurement showed that the enhanced stability of OmicronRBD can be mainly attributed to the S373P mutation.

Next, the unfolding kinetics of the wtRBD and OmicronRBD were determined by stretching the proteins at different velocities during a force loading rate experiment^35–37^. In force spectroscopy measurements, the application of external mechanical force lowers the activation energy of protein unfolding (Fig. 4d). Thus, the unfolding force is proportional to the logarithm of the loading rate, which describes the effect of the force applied on the protein over time. As expected, the plots showed a linear relationship (Figs. 4e, S13). From the fit, we estimated the unfolding rate (*k*_off_) value and the length scale of the energy barrier (Δx) value. The *k*_off_ was 0.59±0.43 s^−1^ (value and fitting error) for the wtRBD and 0.12±0.12 s^−1^ for the OmicronRBD. The *k*_off_ of WT(S373P) decreased to a similar value as that of OmicronRBD, while WT(S371L) remained mostly unchanged. The kinetic results also showed a stabilized OmicronRBD with a lower off rate than the wtRBD, and the mutation S373P plays an important role in the increment of mechanical stability.

Finally, we performed RBD/ACE2 unbinding experiment to study the binding ability of RBD to ACE2, which is the primary biological function of RBD. RBDs and ACE2 were ligated on the AFM tip and glass surface using the previous AEP-based enzymatic method, respectively (Fig. 5a). The AFM tip approached the surface, leading to the complex formation. Then the tip retracted, and the complex unbound upon stretching. The corresponding force-extension curve showed a single peak (Fig. 5b) from the unbinding event between ACE2 and RBD. Then, the tip moves to another position and repeats this cycle thousands of times. We then counted their unbinding events from ten independent experiments. It is found that the binding probability of OmicronRBD is higher than wtRBD (14.6% vs.7.3%, Fig. 5c).

**Figure 5.**
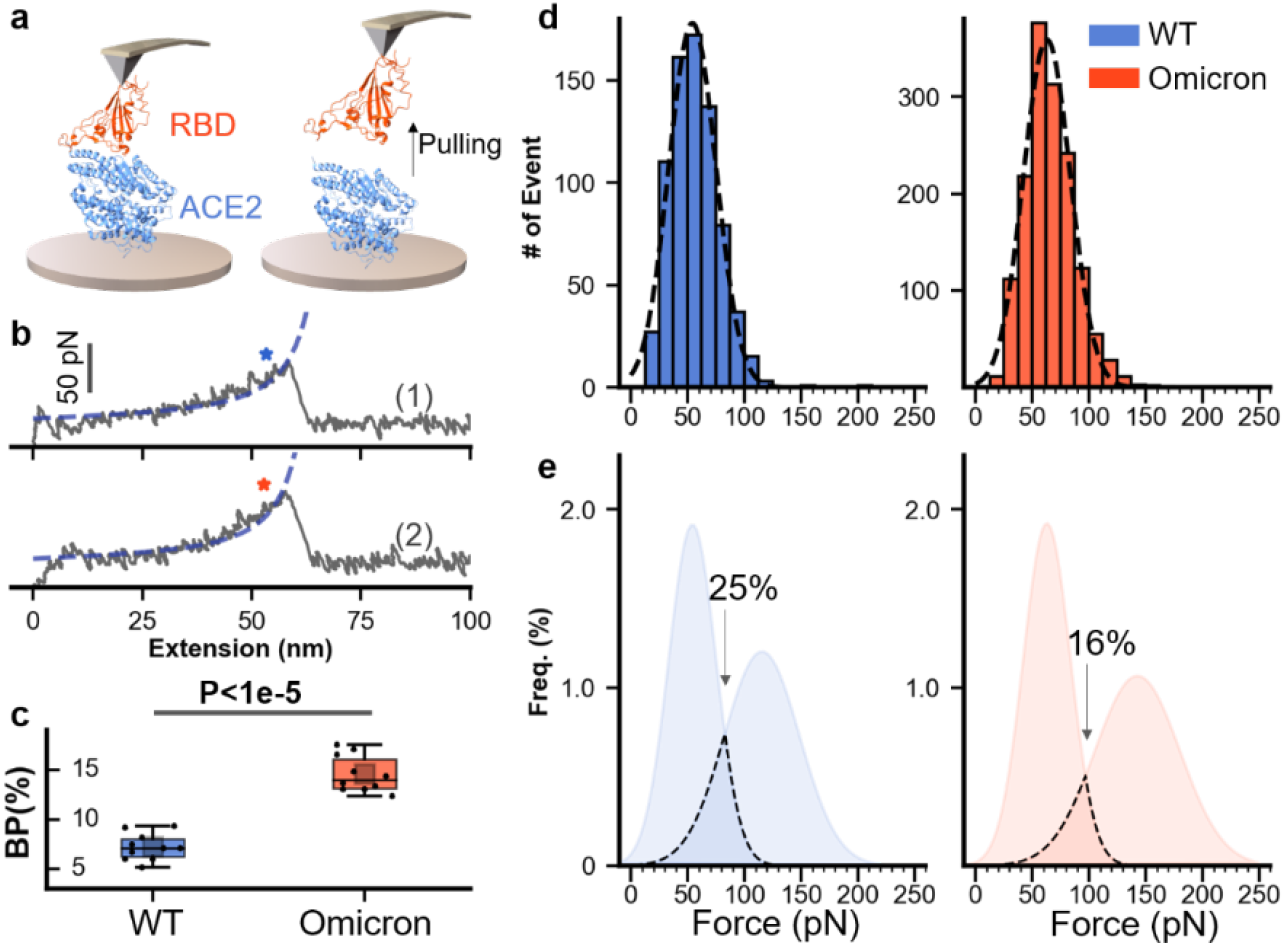
AFM-SMFS unbinding experiment of RBD-ACE2 complex. **a**, Schematic of measurement of the unbinding between RBDs and ACE2 using SMFS. RBD and ACE2 are immobilized to the AFM tip and substrate, respectively. **b**, Representative force-extension curves show the unbinding peak from RBD-ACE2 interaction. WT is colored blue (top), and Omicron is orange (bottom). **c**, Box plot of the binding probabilities shows a higher value for the Omicron than WT (14.6% vs. 7.3%). One data point belongs to the BP from one experiment (ten in total). The box indicates the 25th and 75th percentiles. P values were determined by a two-sample t-test. **d**, The unbinding force distribution of the RBDs-ACE2 complex. Dashed lines represent the gaussian fit of the result. **e**, The merged results of unfolding and unbinding for WT and Omicron RBD. Overlap of the force distribution is observed for both WT (25%) and Omicron (16%) under the experimental condition.

In addition, the average rupture force was 54 ± 21 pN for wtRBD-ACE2 interaction and 63 ± 21 pN for OmicronRBD-ACE2 interaction, determined from the force distribution (Fig. 5d). Finally, we fit the distribution by the gaussian function and merged the unfolding force of RBD and unbinding force (Fig. 5e). An overlapped region between two distribution represents. The overlapped area is 25% for wtRBD and 16% for OmicronRBD, implying that the enhanced mechanical stability of RBD may contribute to the protection of protein structure from unbinding events under our experimental condition.

## Discussion

Coronaviruses are large, single-stranded RNA viruses evolving with a remarkable mutation rate, as evidenced by the transmission of several VOCs of SARS-CoV-2 in only the past two years. In addition to neutral mutations, the effect of many accumulated mutations has been revealed. In this study, we combined experimental and computational approaches to study the mechanical stability of the OmicronRBD and determine the effect of mutation S373P in the β-core region. First, the AFM measurements showed an unfolding force for that OmicronRBD that was 20% higher than its characteristic unfolding. To the best of our knowledge, this is the first experimental work to show higher mechanical stability of the OmicronRBD quantitatively. Here, the binding between RBD and ACE2 as the host-pathogen interaction is a classic protein-protein interaction. It is established that the non-covalent interactions between the two protein contact surfaces are the essential determinant for their binding strength. Here, we showed that the (mechanical) stability of one protein itself could promote its binding ability to its partner.

In addition to the experimental results, the computational data provide essential mechanistic insight into the OmicronRBD, and these simulations were essential parts of our work^46,47^. MD simulations uncovered its detailed unfolding pathway and revealed the underlying mechanism of the stability increment, which is due to the increased number of hydrogen bonds caused by the mutations S373P and S371L.

From a structural perspective, two subdomains with different mechanical stability are present in the RBD. The RBM dominates ACE2 binding and receptor specificity. Before Omicron, most accumulating mutations in the RBD occurred in the RBM. In addition, mutations in the other subdomain, the β-core region consisting of a central β- sheet flanked by α-helices, were rare. In fact, there are four disulfide bonds in the RBD, strongly supporting the overall structure of the RBD and its function. To date, no naturally occurring variant with mutations to these cysteine residues has been observed. Intended cysteine mutation results in the dysfunction of the RBD binding to ACE2, highlighting the importance of a stable RBD structure. In this work, we found that a mutation in the β-core domain can fine-tune the stability of the RBD. And there is indeed an overlap between the unfolding force of RBD and the RBD/ACE2 unbinding force. Thus, it is possible that the enhanced mechanical stability of RBD contributes to its function and virus transmission^15,38^. Nevertheless, this should be investigated using viruses at the cellular level and model animals.

In addition to discovering the effect of mutation S373P for SARS-CoV-2, this work revealed a new way that nature used to increase protein’s mechanical stability by using proline mutation. Proline mutation is often used to decrease protein’s stability by disrupting the hydrogen bond networks in β-sheet region. Here, by introducing proline in the loop close to the β-sheet region, more hydrogen bonds are formed and stabilize the region. Thus, a new strategy to design proteins with enhanced stability is demonstrated. In fact, the importance of mechanical force in immunity has been demonstrated, such as tuning T-cell receptor sensitivity to mechanical force for better CAR T-cell therapy^39^. And so many single-molecule studies of protein, DNA, and RNA show the relationship between mechanical force and their folding and function^40–45^. Thus, mechanical stability can be a factor in SARS-CoV-2 mutation selection.

Finally, S373P in the RBD of spike protein is a highly conserved mutation in Omicron. It is found in all Omicron subvariants, including BA.2 (B.1.1.529.2), BA.3 (B.1.1.529.3), BA.4 (B.1.1.529.4), and BA.5 (B.1.1.529.5), and the very recent strains BQ.1, XBB and CH.1.1^48–51^. Mutation S373P is identical, while S371 is mutated to the more hydrophobic residue leucine in BA.1 and phenylalanine in BA.2-5, further demonstrating the importance of this mutation and suggesting its effect on higher transmission. Since we cannot predict future mutations and emerging variants of SARS-CoV-2, it is critical to maintain surveillance of existing variants and mutations and study their effects.

## Materials and method

### Engineering of Recombinant Protein, Experimental Procedure, Data Analysis

All the different RBD constructs are built by standard molecular biology methods. Their expression, purification, and immobilization in the AFM-SMFS system can be found in SI Appendix for details. AFM-SMFS experiments were performed by Nanowizard 4 atomic force microscope, and the details can be found in SI Appendix. MD simulations were used to reveal the underlying molecular mechanism, and the detail is also provided in SI Appendix.

## Acknowledgments

The numerical calculations in this paper have been done on the computing facilities in the High-Performance Computing Center (HPCC) of Nanjing University.The authors thank inspiring discussions with Prof. Thomas T. Perkins and Prof. Michael M. Nash.

## Funding

National Natural Science Foundation of China grant No. 21977047, 22222703 (PZ) Natural Science Foundation of Jiangsu Province No. BK20200058, BK20202004 (PZ)

## Competing interests

The authors declare no competing interests.

## Data and materials availability

All data are available in the main text or the supplementary materials

## Supplementary Materials inclues

Materials and methods; supplementary text, Figures (S1 to S13), References (1–12) and Movies S1 to S4

## SUPPLEMENTARY MATERIALS

### Materials and Methods

#### Protein expression and purification

The genes were ordered from Genscript Inc. The wtRBD and OmicronRBD constructs contain the SAS-CoV-2 spike protein (residues 319-591), followed by a GGGGS linker and an 8XHis tag in pcDNA3.4 modified vector. They were expressed in Expi293 cells with OPM-293 CD05 serum-free medium (OPM Biosciences)(*20*). In addition to these two RBDs, most RBDs (residues 333-528) were constructed as a fused polyprotein Coh-(GB1)_2_-RBD-GB1-NGL for high-precision AFM measurement and thus expressed in *E. coli* BL21(DE3) using pQE80L vector. The *BamH*I*-Bg*lII*-Kpn*I three-restriction enzyme system was used for the stepwise construction of the genes for polyproteins. For protein purification of RBD with His-tag, culture supernatant was passed through a Ni-NTA affinity column (Qiagen). Proteins were further purified by gel filtration (Superdex™ 200 Increase 10/30GL, GE Healthcare).

*Oa*AEP1(C247A) is cysteine 247 to alanine mutant of asparaginyl endoproteases 1 from *oldenlandia affinis*, abbreviated as AEP here (*1*). ELP is the elastin-like polypeptides (*2*). Their expression and purification protocols can be found in references. RBD mutants, including wtRBD(S371L), wtRBD(S373P), OmicronRBD (L371S), and OmicronRBD (P373S), were generated using the QuikChange kit. Their sequences were all verified by direct DNA sequencing.

For the RBDs-ACE2 unbinding experiment, the genes were ordered from GenScript Inc. The RBDs (WT, Omicron) construct contains the SARS-CoV-2 spike protein (residues 319–591), followed by a GGGGS linker and an His8 tag in a pcDNA3.4 modified vector. Their sequences were all verified by direct DNA sequencing (GENERAL BIOL). A C-terminal NGL was added to the RBDs for ligation. The human ACE2 construct contains the ACE2 extracellular domain (residues 19–740) and an Fc region of IgG1 at the C-terminus followed by NGL.

#### AFM-SMFS unfolding experiment

The AFM cantilever/tip made of silicon nitride (MLCT-BIO-DC, Bruker Corp.) was used. The detailed protocol for AFM tip functionalization and protein immobilization on the glass coverslip can be found in references (*30, 3*). In short, the tip and glass coverslip were coated with the amino group by amino-silanization. Then, the maleimide group for cysteine coupling was added on the amino-functionalized surface using the hetero-bifunctional crosslinker sulfosuccinimidyl 4-(N-maleimidomehthyl) cyclohexane-1-carboxylate) (Sulfo-SMCC, Thermo Scientific). Next, the peptide GL-ELP_20_-C or C-ELP_20_-NGL was reacted to the maleimide via the cysteine, respectively. The long ELP_20_ serves as a spacer to avoid non-specific interaction between the tip and the surface as well as a signature for the single-molecule event. Finally, target protein RBDs with C-terminal NGL sequence or GB1-Doc with N-terminal GL sequence can be site-specifically linked to the coverslip or tip by ligase AEP, respectively.

Atomic force microscope (Nanowizard4, JPK) was used to acquire the force-extension curve. The D tip of the MLCT-Bio-DC cantilever was used. Its accurate spring constant was determined by a thermally-induced fluctuation method (*4*). Typically, the tip contacted the protein-immobilized surface for 400 *ms* under an indentation force of 450 pN to ensure a site-specifically interaction. Then, moving the tip up vertically at a constant velocity (1 μm/s, if not specified), the polyprotein unfolded. Then, the tip moved to another place to repeat this cycle several thousands of times. As a result, a force-extension curve was obtained, which was analyzed using JPK data process analysis software.

#### AFM-SMFS unbinding experiment of RBD-ACE2 on surface

FD-based AFM on model surfaces was performed in Tris-HCl buffer (pH7.4, 150 mM NaCl, 50 mM Tris-HCl) at room temperature using functionalized D tip of MLCT-Bio-DC cantilever (Bruker, nominal spring constant of 0.030 N/m and actual spring constants calculated using thermal tune). AFM (Nanowizard4, JPK) operated in the force mapping (contact) mode was used. Areas of 10 × 10 μm were scanned, ramp size set to 350 nm, and set point force of 300 pN with a contact time of 50 ms, with a resolution of 32 × 32 pixels.

#### Bell-Evans model to extract kinetics

The RBD-ACE2 complex dissociation in the AFM experiment is a non-equilibrium process that can be modeled as an all-or-none two-state process with force-dependent rate constant *k*(F). The rate constant can be described by Bell-Evans’ model (*34*):

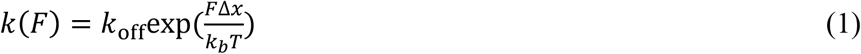

*k*(*F*) is the protein unfolding rate under a particular force *F*, *k*_off_ is the unfolding rate constant under zero force, Δx is the distance between the unfolded state and the transition state. For the dynamic force spectroscopy measurements, the slope *aa* of the force−extension curves immediately before the unfolding event (~2 nm) was first determined to obtain the average loading rate (*rr* = *aaaa*, where *aa* is the velocity). The Bell-Evans model was used to fit all the data (1), yielding the spontaneous unfolding rate, and the distance from the folded state to the transition state with the following equation:

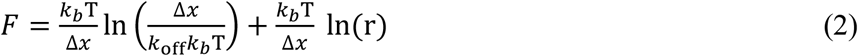

Four different pulling velocities, 0.2 μm/s, 0.4 μm/s, 1 μm/s, and 4 μm/s were used. The relationship between the most probable unfolding force and loading rate was obtained on a log scale, which is fitted by a linear line as equation (2).

#### MD simulation for RBD unfolding

To reveal the unfolding pathway of RBD under mechanical load, we performed steered molecular dynamics (SMD) simulations using NAMD(*5, 6*). Simulation systems were prepared using CHARMM-GUI (*7*). The structures of the RBDs were prepared following established protocols. For the WT, the structure had been solved by Electron Microscopy at 2.60 Å resolution and is available at the protein data bank (PDB ID: 6zge). The Omicron had also been solved by Electron Microscopy, at 3.5 Å resolution, and is available at the protein data bank (PDB ID: 7tl1). After extracting residues 333 to 528 in part A both of them, the RBDs were solvated and the net charge of the proteins were neutralized using a 150 mM salt concentration of sodium chloride. Disulfide bonds and the glycans at N343 were included following the literature information (*8*). SMD simulations were performed employing the NAMD molecular dynamics package. The CHARMM36 force field along with the TIP3P water model was used to describe all systems. The simulations were performed assuming periodic boundary conditions in the NpT ensemble with temperature maintained at 298 K using Langevin dynamics for pressure, kept at 1 bar, and temperature coupling (*9*). Before the MD simulations, all the systems were submitted to an energy minimization protocol for 5,000 steps. MD simulations with position restraints in the protein backbone atoms were performed for 1.0 *ns* and served to pre-equilibrate systems before the 10 *ns* equilibrium MD runs. To characterize the unfolding pathway of RBDs, ten times of SMD simulations with constant stretching velocity employed a pulling speed of 5.0 Å/ns and a harmonic constraint force of 7.0 kcal/mol/Å was applied for 30.0 ns. In this step, SMD were employed by harmonically restraining the position of the C-terminus and pulling on the N-terminus of the RBDs (WT or Omicron). Each system was run 10 times. Simulation force-extension traces were analyzed analogously to experimental data. Data were analyzed by python-based Jupyter notebooks. Jarzynski’s equality is applied to potential mean force (PMF) from SMD simulations (*10*). We also performed 60 times of SMD simulations with a pulling speed of 20 Å/ns to calculate the PMF.

MD simulations were performed utilizing the GROMACS 2021 package. And all simulation systems, including the mutation (G339D, S371L, S373P, or S375F), is established with similar procedures mentioned above including energy minimization and pre-equilibrium. All the input files were generated from CHARMM-GUI. The system’s environment changed at 298 K (NVT ensemble) and subsequently at 298 K and 1 bar (NPT ensemble). We performed 300 ns MD simulations 10 times. And the number of hydrogen bonds formed between β2/β10 and β1/β5 (residue 353 to 363, 394 to 400, 523 to 526) was analyzed by the plug-in program *hbond* in GROMACS(*11*).

The cutoff parameters of the hydrogen bond analysis program were defaulted values (3.5 Å and 30°). It means that when the distance between donor D and acceptor A was shorter than 3.5 Å, as well as the bond angle H–D is smaller than 30.0°, it is regarded as a hydrogen bond. Hydrogen bond formation for every residue between β2/β10 and β1/β5 is also generated by the same plug-in program, executed by a Python script. The distance between the Cα atoms of the 373 and 343 residues during the MD simulations of WT and Omicron was calculated by another plug-in program *distance* and organized by a python script to form a scatter plot with the number of hydrogen bonds. Then we use the pearsonr function in the scipy module to evaluate the correlation between the number of hydrogen bonds and the distance between residue 371 and 343 (*12*). RMSF and energy analysis were performed using gromacs built-in programs *rmsf* and *energy*

## Supplementary Text

### Protein sequences

#### His6-Coh-GB1-GB1-RBD**(XX)**-GB1-NGL

MRGSHHHHHHGSMGTALTDRGMTYDLDPKDGSSAATKPVLEVTKKVFDTAADAAGQTVTVEFKVSGAEGKYATTGYHIYWDERLEVVATKTGAYAKKGAALEDSSLAKAENNGNGVFVASGADDDFGADGVMWTVELKVPADAKAGDVYPIDVAYQWDPSKGDLFTDNKDSAQGKLMQAYFFTQGIKSSSNPSTDEYLVKANATYADGYIAIKAGEPRSMDTYKLILNGKTLKGETTTEAVDAATAEKVFKQYANDNGVDGEWTYDDATKTFTGTERSMDTYKLILNGKTLKGETTTEAVDAATAEK VFKQYANDNGVDGEWTYDDATKTFTGTERSXXRSMDTYKLILNGKTLKGET TTEAVDAATAEKVFKQYANDNGVDGEWTYDDATKTFTVTERSNGL

#### XX: **WT**

TNLCPFGEVFNATRFASVYAWNRKRISNCVADYSVLYNSASFSTFKCYGVSPTKLNDLCFTNVYADSFVIRGDEVRQIAPGQTGKIADYNYKLPDDFTGCVIAWNSNNLDSKVGGNYNYLYRLFRKSNLKPFERDISTEIYQAGSTPCNGVEGFNCYFPLQSYGFQPTNGVGYQPYRVVVLSFELLHAPATVCGPK

#### XX: **WT(C4^−^)**

TNLCPFGEVFNATRFASVYAWNRKRISNCVADYSVLYNSASFSTFKAYGVSPTKLNDLAFTNVYADSFVIRGDEVRQIAPGQTGKIADYNYKLPDDFTGAVIAWNSNNLDSKVGGNYNYLYRLFRKSNLKPFERDISTEIYQAGSTPCNGVEGFNCYFPLQSYGFQPTNGVGYQPYRVVVLSFELLHAPATVAGPK

#### XX: **WT(S371L)**

TNLCPFGEVFNATRFASVYAWNRKRISNCVADYSVLYNLASFSTFKCYGVSPTKLNDLCFTNVYADSFVIRGDEVRQIAPGQTGKIADYNYKLPDDFTGCVIAWNSNNLDSKVGGNYNYLYRLFRKSNLKPFERDISTEIYQAGSTPCNGVEGFNCYFPLQSYGFQPTNGVGYQPYRVVVLSFELLHAPATVCGPK

#### XX: **WT(S373P)**

TNLCPFGEVFNATRFASVYAWNRKRISNCVADYSVLYNSAPFSTFKCYGVSPTKLNDLCFTNVYADSFVIRGDEVRQIAPGQTGKIADYNYKLPDDFTGCVIAWNSNNLDSKVGGNYNYLYRLFRKSNLKPFERDISTEIYQAGSTPCNGVEGFNCYFPLQSYGFQPTNGVGYQPYRVVVLSFELLHAPATVCGPK

#### XX: **Omicron**

TNLCPFDEVFNATRFASVYAWNRKRISNCVADYSVLYNLAPFFTFKCYGVSPTKLNDLCFTNVYADSFVIRGDEVRQIAPGQTGNIADYNYKLPDDFTGCVIAWNSNKLDSKVSGNYNYLYRLFRKSNLKPFERDISTEIYQAGNKPCNGVAGFNCYFPLRSYSFRPTYGVGHQPYRVVVLSFELLHAPATVCGPK

#### XX: **Omicron(4C-)**

TNLCPFDEVFNATRFASVYAWNRKRISNCVADYSVLYNLAPFFTFKAYGVSPTKLNDLAFTNVYADSFVIRGDEVRQIAPGQTGNIADYNYKLPDDFTGAVIAWNSNKLDSKVSGNYNYLYRLFRKSNLKPFERDISTEIYQAGNKPCNGVAGFNCYFPLRSYSFRPTYGVGHQPYRVVVLSFELLHAPATVAGPK

#### XX: **Omicron(L371S)**

TNLCPFDEVFNATRFASVYAWNRKRISNCVADYSVLYNSAPFFTFKCYGVSPTKLNDLCFTNVYADSFVIRGDEVRQIAPGQTGNIADYNYKLPDDFTGCVIAWNSNKLDSKVSGNYNYLYRLFRKSNLKPFERDISTEIYQAGNKPCNGVAGFNCYFPLRSYSFRPTYGVGHQPYRVVVLSFELLHAPATVCGPK

#### XX: **Omicron(P373S)**

TNLCPFDEVFNATRFASVYAWNRKRISNCVADYSVLYNLASFFTFKCYGVSPTKLNDLCFTNVYADSFVIRGDEVRQIAPGQTGNIADYNYKLPDDFTGCVIAWNSNKLDSKVSGNYNYLYRLFRKSNLKPFERDISTEIYQAGNKPCNGVAGFNCYFPLRSYSFRPTYGVGHQPYRVVVLSFELLHAPATVCGPK

---

Movies S1 to S4

S1: Movie of Steering molecular dynamics (SMD) simulations of wtRBD

S2: Movie of Steering molecular dynamics (SMD) simulations of OmicronRBD S3: Movie of MD simulations to determine the number of H-bonds of the wtRBD

S4: Movie of MD simulations to determine the number of H-bonds of the OmicronRBD

**Figure S1.**
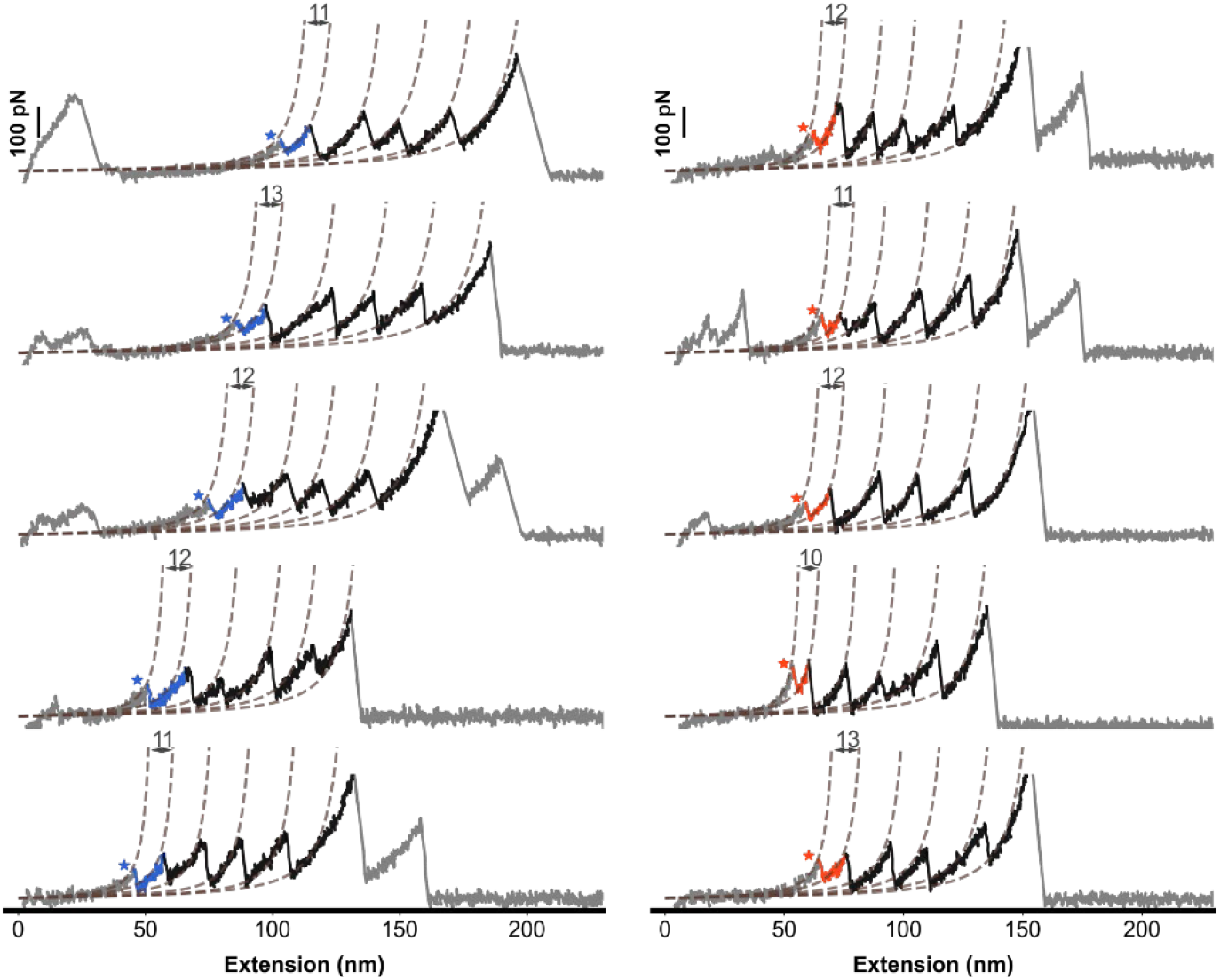
More representative curves of RBDs. Representative curves showing a series of sawtooth-like peaks from the unfolding of the RBD with a ΔLc of ~11 nm and marker protein GB1 with a ΔLc of ~18 nm. The Omicron and WT variants are colored orange and blue, respectively.

**Figure S2.**
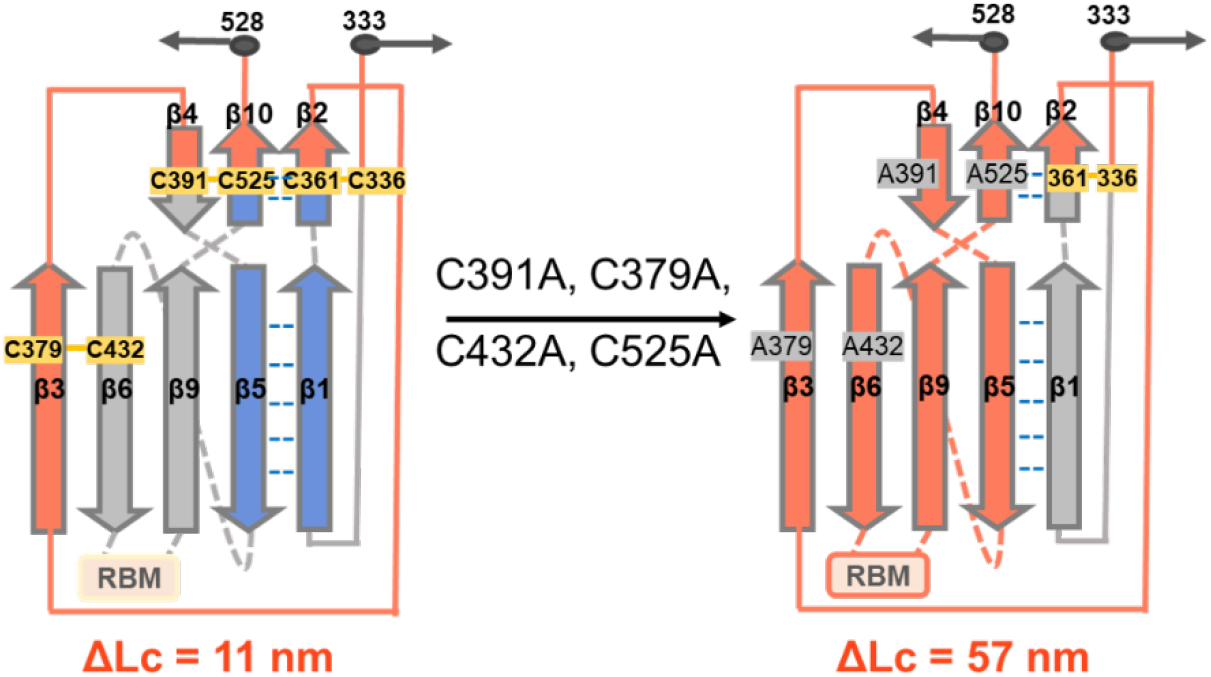
The unfolded fragments of RBD and RBD (4C^−^) upon unfolding. . The comparison between RBD and Cys-delete RBD mutant cartoon structure indicates that a larger ΔLc (57 nm) upon unfolding is expected due to the release of a more extensible protein structure (colored in orange) upon the deletion of two disulfide bonds.

**Figure S3.**
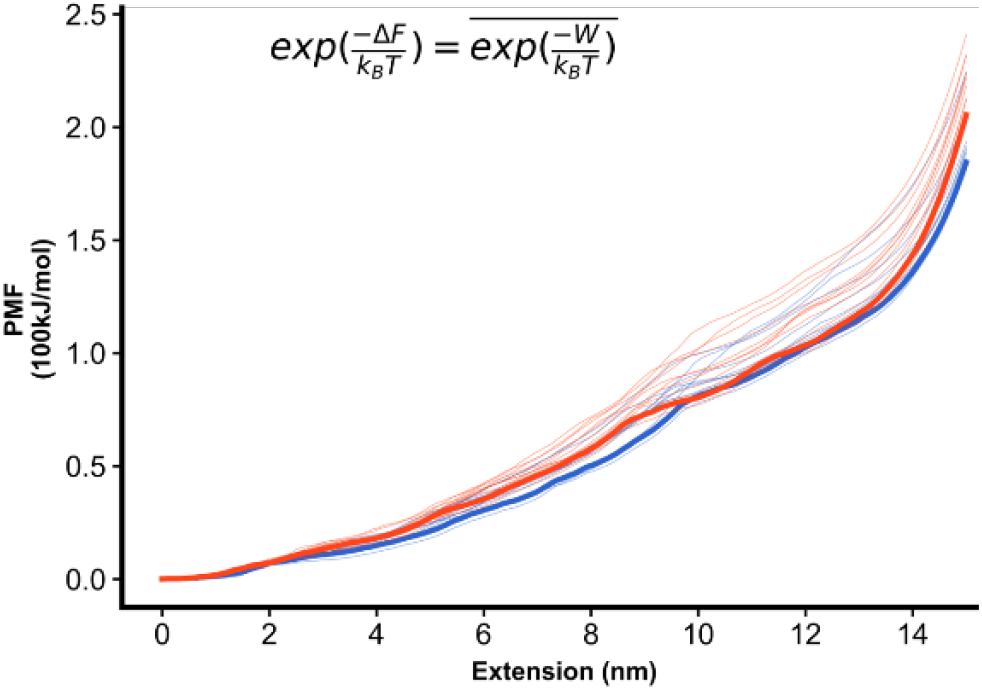
PMF calculation shows that Omicron is more stable than WT. PMF was calculated from irreversible pulling (*v* =5 Å/ns) of 10 SMD trajectories. Light color lines represent the work done in the simulations.

**Figure S4.**
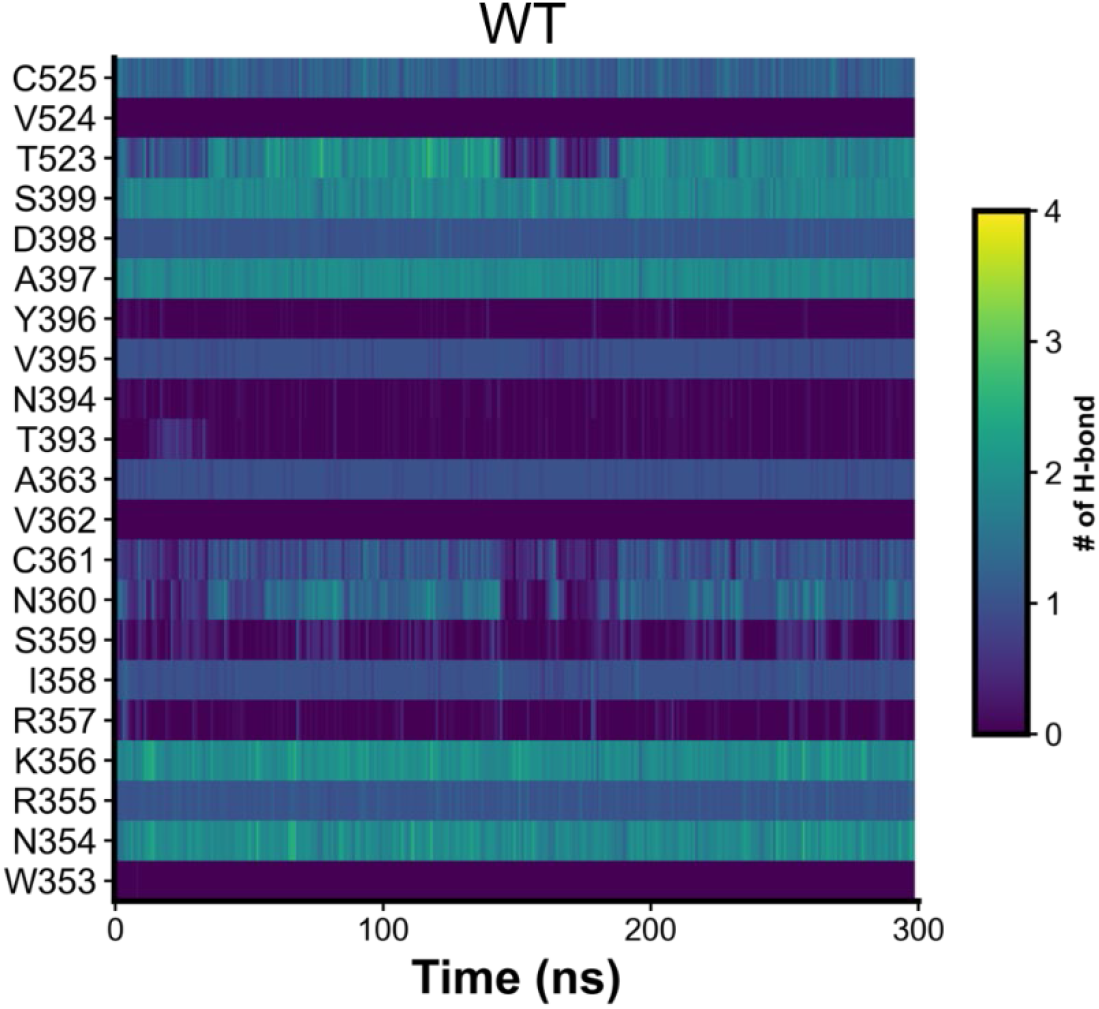
The evolution of H-bond number for each residue involved in β2/β10 and β1/β5 during the MD simulations for wtRBD.

**Figure S5.**
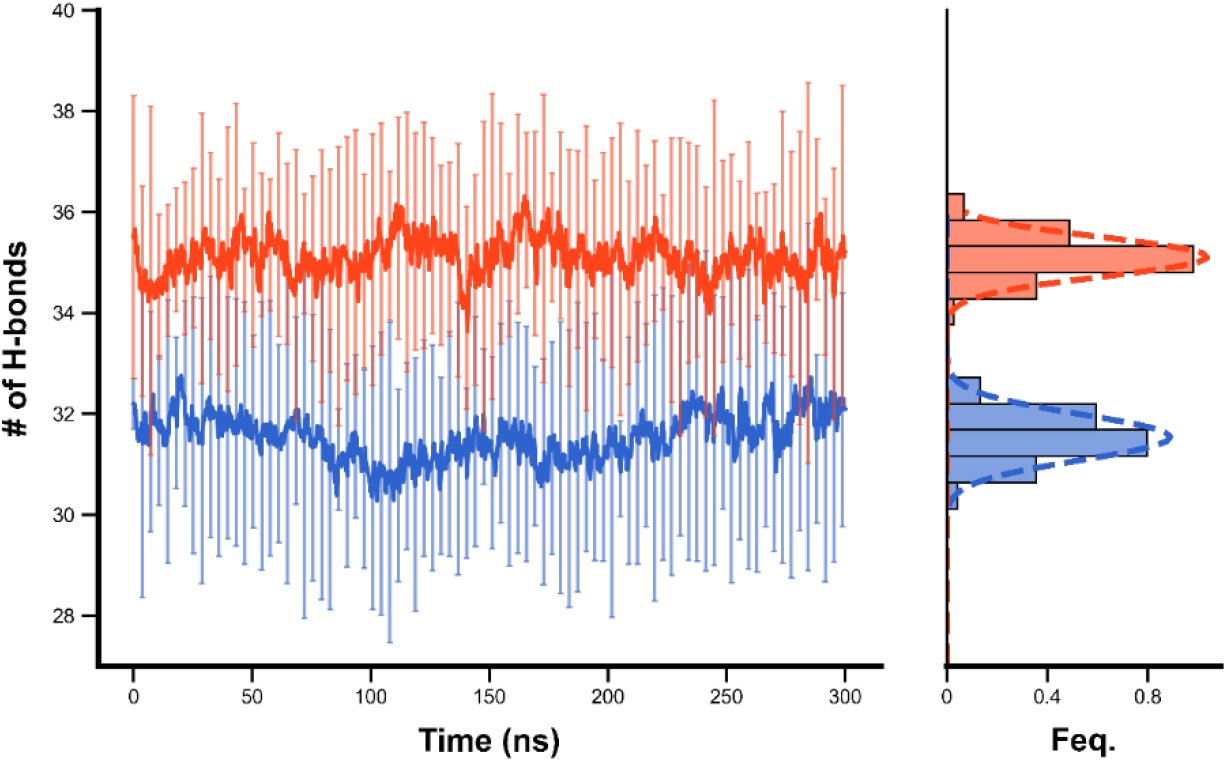
Overall MD simulations during 300 ns. It shows the average number of hydrogen bonds formed in the β-core region. The average number of hydrogen bonds formed by OmicronRBD in the β-core region is still larger than that of wtRBD.

**Figure S6.**
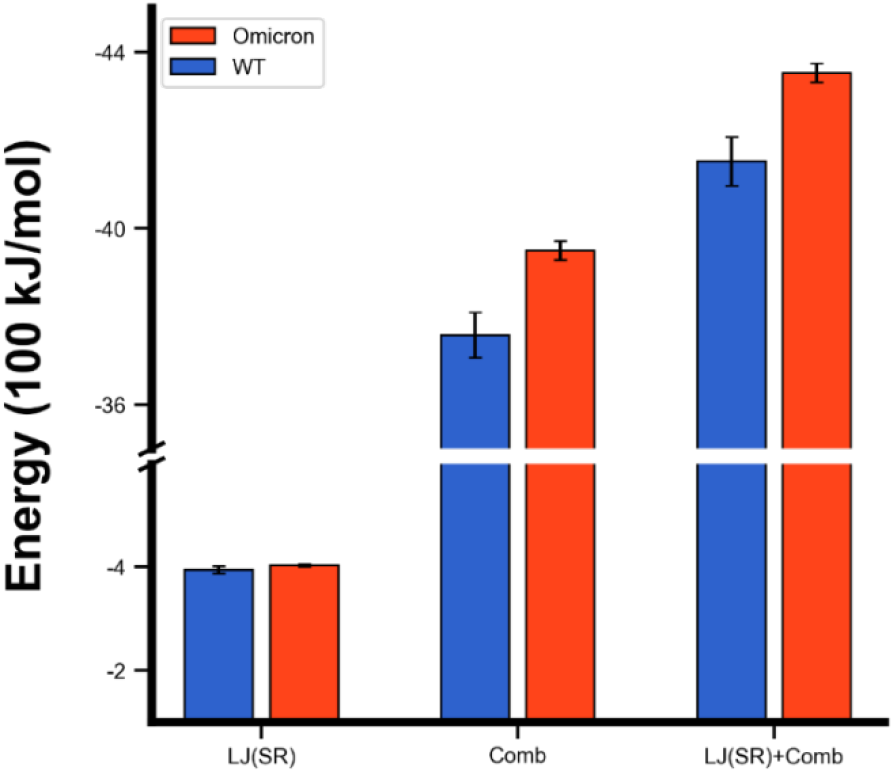
Energy analysis of β-strands during 10 times MD simulations. The coulomb interaction of Omicron RBD is stronger than wtRBD. Thus, the Coulomb interaction might play an important role in the stability of the β-strands.

**Figure S7.**
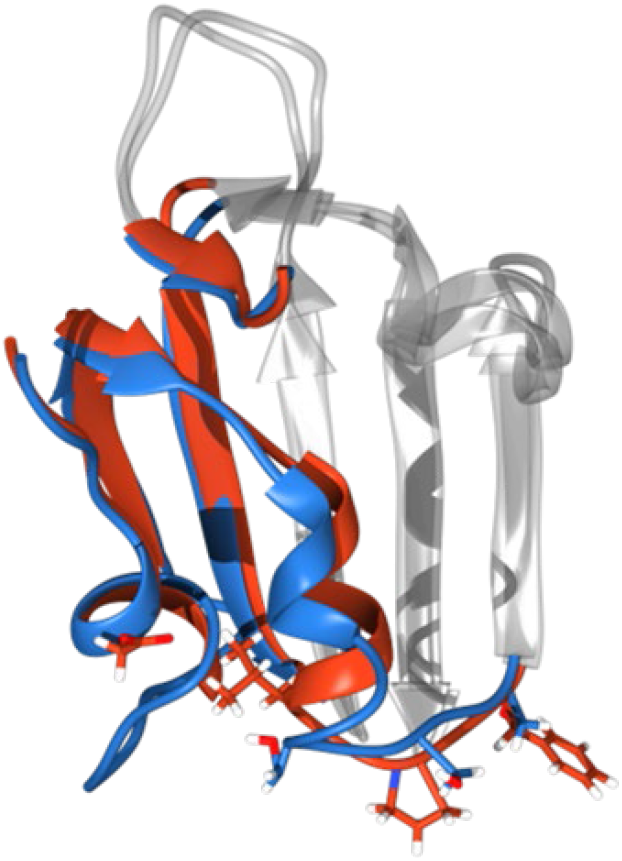
Overlap of the Cryo-EM structure of Omicron (red) and WT (blue). A slight conformational change closing to the β-core region of RBD is observed between Omicron and WT. The four mutations are depicted.

**Figure S8.**
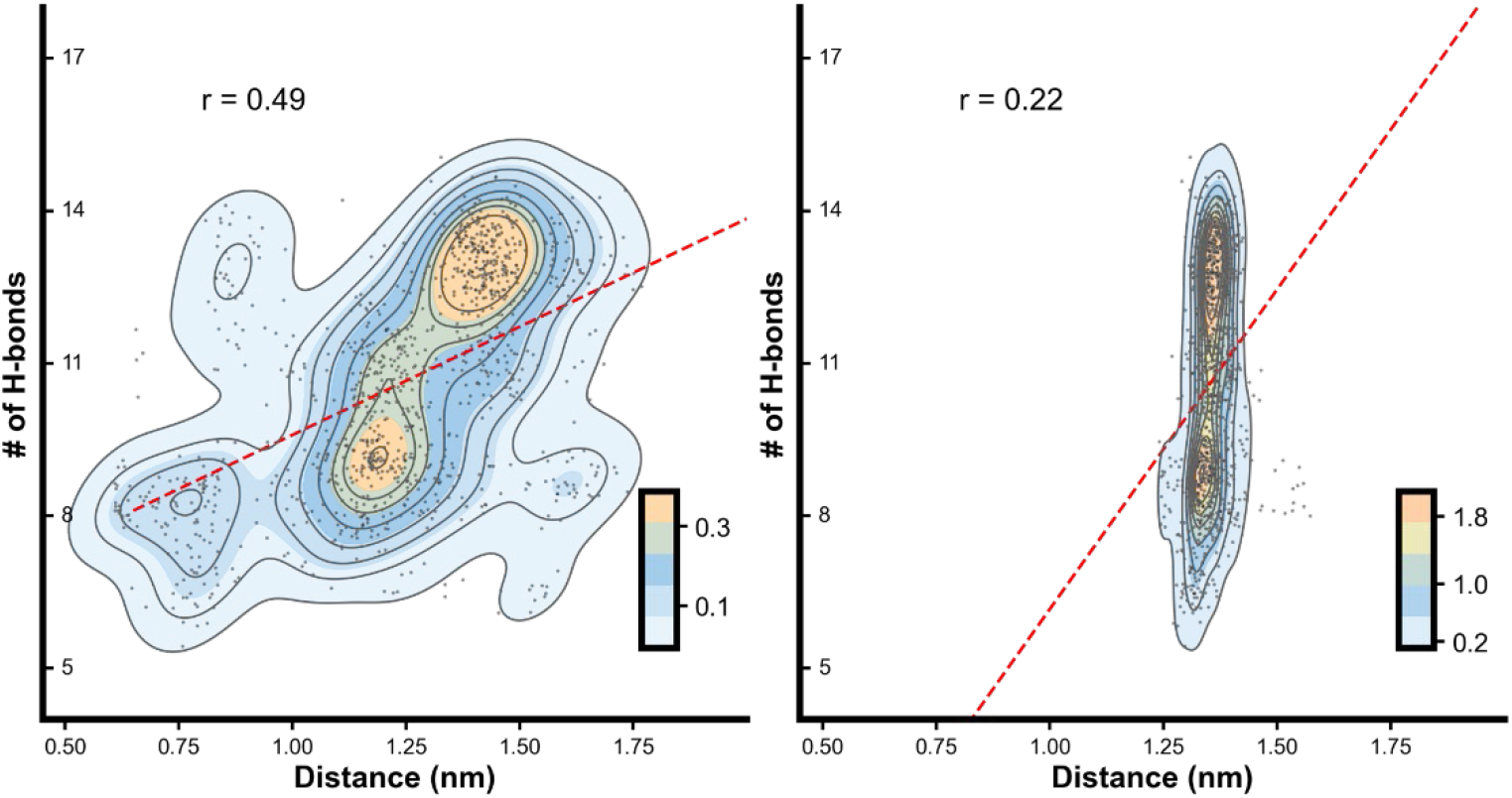
The relationship between the distances and the number of hydrogen bonds for different residues. The relationship between the distances of residues 371(left) / 375(right) to 432 and the number of hydrogen bonds is week.

**Figure S9.**
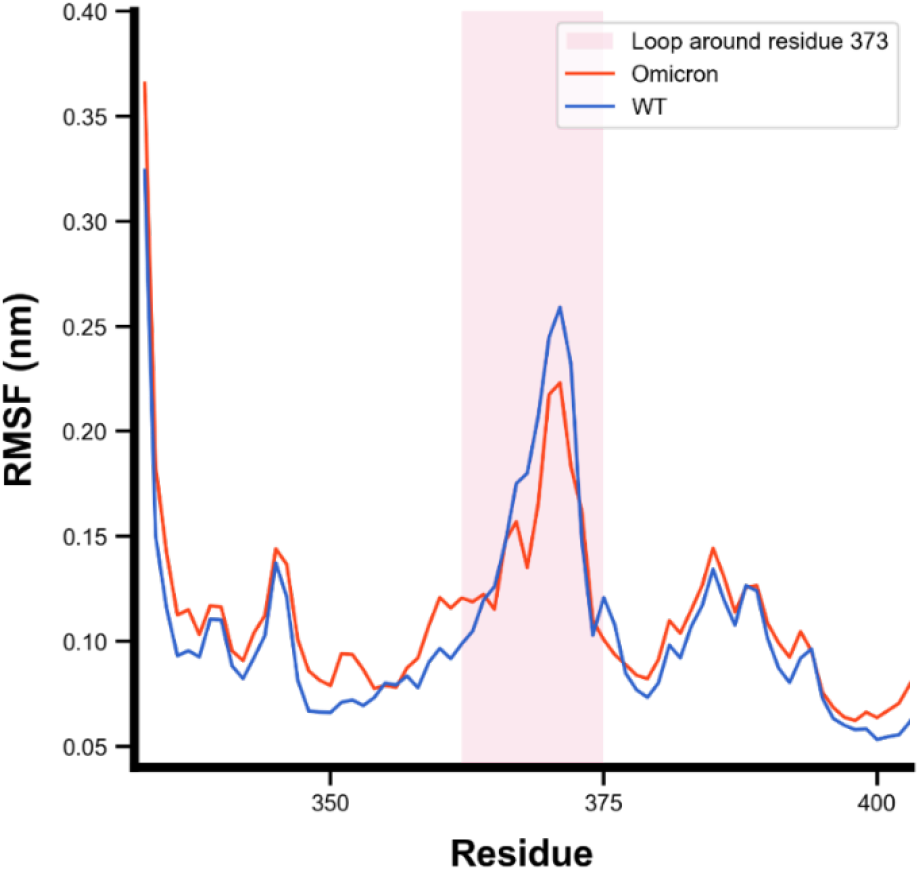
The average RMSF value of 10 times MD simulations. The RMSF of helix around residue 373 in OmicronRBD was lower than wtRBD, suggesting that the mutation of OmicronRBD could enhance the stability of this region and change the movement of specific residues.

**Figure S10.**
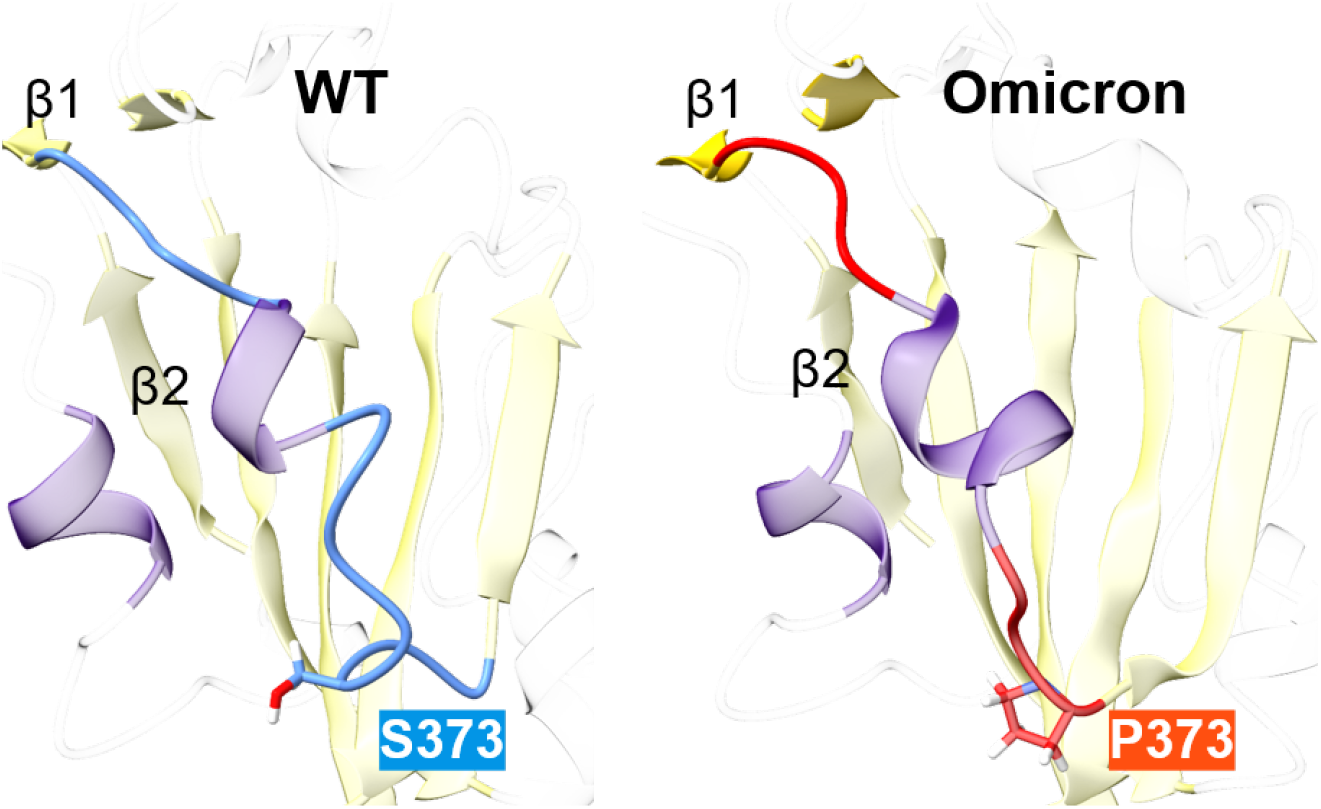
A focus view of peptide chain around residue 373. The rigid proline 373 in OmicronRBD might open the loop between the nearby helix and β-strand, so that it can maintain a relatively large distance from the helix.

**Figure S11.**
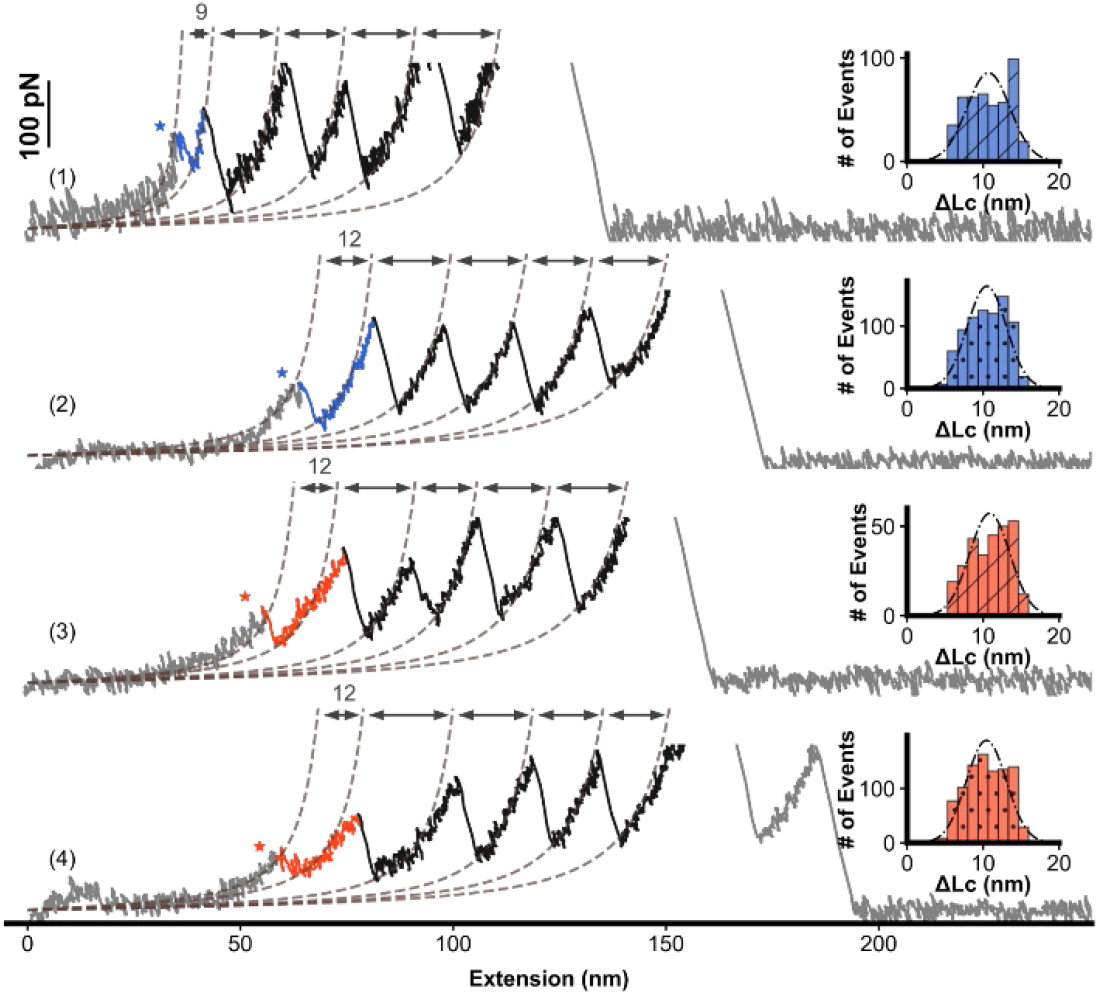
Representative curves of single mutation of residue 371 or 373 on RBD. Force-extension curves of RBDs of WT(S371L), WT(S373P), Omicron(L371S), and Omicron(P373S) showed the stepwise unfolding of the polyprotein, including the RBD, respectively. The histogram of ΔLc of RBD is shown in the inset.

**Figure S12.**
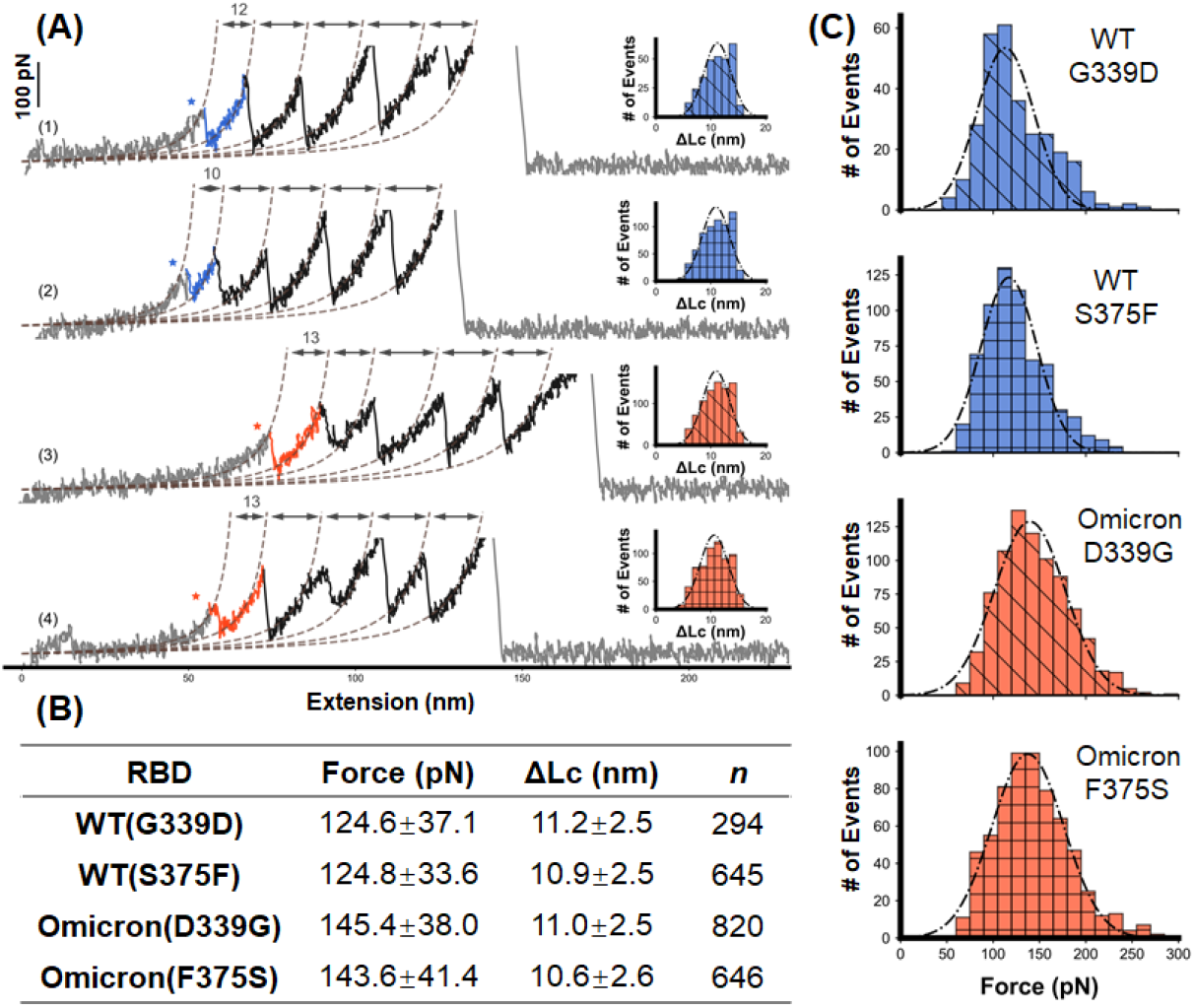
AFM unfolding results of single mutation of residue 339 or 375 on RBD. **A)** Force-extension curves of RBDs of WT(G339D), WT(S375F), Omicron(D339G), and Omicron(F375S) showed the stepwise unfolding of the polyprotein, including the RBD, respectively. The histogram of ΔLc of RBD is shown in the inset. **B-C)** AFM unfolding results of the four mutants are shown in details.

**Figure S13.**
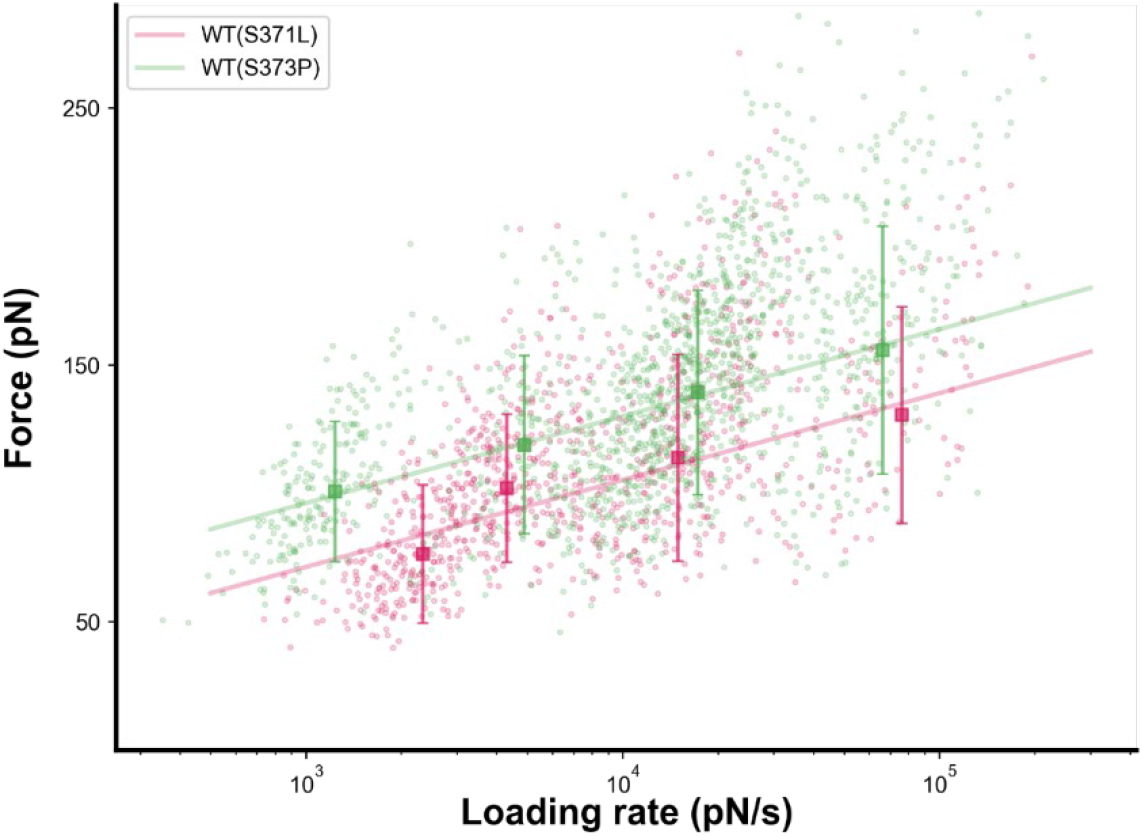
Dynamic force spectrum of the unfolding of RBDs. The unfolding forces of the WT(S371L) and WT(S373P) show a linear relationship with the logarithm of the loading rate. The *k*_off_ was 0.53±0.79s^−1^ for the WT(S371L) and 0.10±0.04 s^−1^ for the WT(S373P). The WT(S371L) and WT(S373P) are colored pink and green, respectively.

## Notes

### Competing Interest Statement

The authors have declared no competing interest.

### Summary of Updates

the unbinding measurement have been included.

